# The role of container periphyton as an oviposition attractant for female *Aedes aegypti*

**DOI:** 10.64898/2025.12.05.692399

**Authors:** Luisa M. Otero-Lopez, Alexandra J. Bauer, Eric P. Caragata, Roberto Barrera

## Abstract

**Background:** *Aedes aegypti* mosquitoes use diverse chemosensory cues to select an oviposition site. These cues are thought to provide information on factors such as food availability, predators, pathogens, and competitors. Most research on the stimuli attracting female *Ae. aegypti* to oviposit in natural or artificial containers has focused on the properties of the vessel and water, such as size, color, water quality, concentration of dissolved organic matter, and microorganisms. One understudied aspect of oviposition containers is periphyton, the complex community of microorganisms that can grow on the inner walls of containers where gravid *Ae. aegypti* females lay their eggs. Periphyton can potentially act as an indicator of the container’s suitability for females looking for an oviposition site, and as a source of food for future mosquito larvae. We were interested in understanding if container periphyton attracts gravid *Ae*. *aegypti* females for oviposition and to assess the microbial composition of the periphyton community.

**Methods:** To answer these questions, we conducted a series of attraction and oviposition assays and collected samples from the periphyton used in these assays for 16s rRNA fingerprinting.

**Results:** We demonstrated that periphyton promotes oviposition by *Ae. aegypti.* Results from 16s rRNA fingerprinting of periphyton specimens revealed that these communities exhibit high complexity and contain microbial taxa from several classes, including Cyanobacteria, Proteobacteria, Anaerolineae, and Cytophagia.

**Conclusions:** Understanding the chemosensory interactions between mosquitoes and periphyton involved in egg laying could prompt the development and formulation of novel attractants useful in mosquito control.

## Background

The adaptation of *Aedes aegypti* (L.) to live within human cities and lay eggs in a diverse array of natural and artificial water-collecting containers has been a major factor in its status as a key vector of the pathogens that cause diseases like dengue, yellow fever, Chikungunya, and Zika fever. The success of this species depends on complex chemosensory biology that facilitates the detection of vertebrate hosts for blood feeding and aquatic habitats for egg laying. Various physical and chemical properties of water collecting vessels define the suitability of these sites for egg laying by mosquitoes. Female mosquitoes can identify and assess oviposition site suitability at variable distances, depending on the cues available. These cues influence their decision to lay eggs in a particular container and ultimately impact the reproductive success of the mosquito population [1].

There are many different types of cues associated with the choice of the place to lay eggs. This includes visual indicators such as the color, volume and diameter of the container [2, 3], and whether the container is in shade [4]. Olfactory and microbial factors including the presence of key microorganisms and the kairomones they release [5, 6], and the concentration of microorganisms [7] are also important. There are also visual and olfactory cues associated with the presence of conspecific eggs [8, 9], the presence and density of intraspecific and interspecific larvae and pupae [10, 11], whether or not those juveniles are starved [12], the presence of parasites [13], and the presence of aquatic predators, including copepods [14]. Water temperature and the texture of the container’s internal surface can also play a role [11, 15].

Another important characteristic of these aquatic environments is the presence of plants and plant-derived compounds. Different plant infusions have demonstrated to be attractive to *Ae. aegypti* females looking to oviposit [16, 17, 18, 19, 20]. Decaying organic matter shapes the abundance and diversity of microorganisms that mosquito larvae might use as a food source [21]. Microbial diversity is high in both natural and artificial containers commonly utilized as larval sources by *Ae. aegypti* [22]. Members of these microbial communities produce an array of secondary metabolites, some of which have odorant properties that are attractive to female mosquitoes and inform about food availability and habitat quality of a prospective oviposition site [7]. Taking advantage of these attractant properties, several different plant infusions, prepared to standardized concentrations, have been utilized in mosquito control programs [23, 24, 25].

One aspect of the aquatic habitats of *Ae. aegypti* that has not been studied is whether the community of microorganisms that inhabits the container wall below and right above the water line, plays a role in attracting gravid *Ae. aegypti* to lay eggs. As water levels in a container decrease, the periphyton (defined as the community of bacteria, algae, and fungi along with organic and inorganic detritus that live on surfaces beneath the water) [26] becomes exposed to air. These periphyton communities are dynamic and can change over time or due to variation in environmental conditions [27, 28, 29, 30]. We hypothesized that the presence of periphyton in a container, or the presence of periphyton-exposed water would promote oviposition from gravid *Ae. aegypti* females. We tested this premise through a series of oviposition choice assays involving the presence or absence of periphyton. We then characterized the richness and diversity of periphyton communities that formed on plastic buckets, a common aquatic habitat for *Aedes aegypti* in urban environments. Our findings provide vital insight into the stability and long-term viability of containers as habitats for mosquito larvae, while potential attractive properties of periphyton could prove useful for mosquito control in the development of baits targeting gravid female mosquitoes.

## Methods

### Oviposition choice experiments

We conducted a set of 2 experiments to determine if containers where we grew periphyton were more attractive to oviposition by *Ae. aegypti* than containers without periphyton and dechlorinated water. Additionally, to understand if the water of containers where we grew periphyton was the cause of attraction, we compared oviposition by *Ae. aegypti* in containers with periphyton and water versus containers without periphyton and periphyton water.

The first experiment (experiment 1) compared oviposition in buckets where periphyton had been allowed to grow against oviposition in control buckets where periphyton had never grown. Water was drained from 3 periphyton buckets, and replaced with 2.5L of fresh, dechlorinated water. An identical volume of dechlorinated water was then added to three control buckets. Thus, there were 3 buckets with periphyton and 2.5 liters of dechlorinated clean water, and 3 control buckets with no periphyton on the walls but 2.5 liters of dechlorinated water. This experimental design was repeated five times.

The second experiment (experiment 2) compared the attractiveness of buckets where periphyton had been allowed to grow against control buckets containing periphyton-exposed water but without an established periphyton layer on the wall of the bucket. For this, we removed 2.5L of water from 3 periphyton buckets and placed it into each of three control buckets. Thus, there were3 buckets with a visible layer of periphyton and 2.5 liters of periphyton water, and 3 control buckets with no periphyton on the walls but 2.5 litters of periphyton water. This experimental design was repeated six times. All oviposition choice experiments were performed between July 21, 2022, and August 26, 2022.

In each experimental replicate, the 6 buckets (three per treatment) were placed inside a semi-field cage (L – 3.35m, W – 1.83m, H - 3.6m) located at the Centers for Disease Control and Prevention (CDC) - Dengue Branch campus. Due to its position, the cage was mostly shaded during the day. Average cage temperature during experiments was 26.93°C (± 2.49, s.d.), while the average relative humidity was 72.13% (± 8.24, s.d.).

Three days prior to each experimental replicate, a cage containing the Patillas strain of *Ae. aegypti* mosquitoes was offered a blood meal using a sausage filled with warm pig’s blood from a local butcher, with 75 mL of anticoagulant added per 425 mL of blood. After feeding, 80 female mosquitoes with blood-filled abdomens were carefully transferred to a separate cage and provided with cotton balls soaked in 10% sucrose.

On the day of the experiment (72 hours post-blood feeding), at 2:00 pm, the cage containing the 80 females (31 cm x 31 cm x 31 cm, metallic) was placed at the center of the semi-field cage. Six equidistant points were marked at 60° intervals along the circle center to serve as bucket locations. Six buckets (three control and three periphyton) were randomly assigned to these points using the random.org list generator (Fig. 1A). At 3:00 pm, the cage was opened, allowing the mosquitoes to enter the semi-field cage. At 11:00 am the following morning, all released mosquitoes were collected using an entomological net, and any carcasses found on the floor were also collected. All specimens were frozen and counted to confirm that all 80 mosquitoes had been removed from the cage.

**Fig. 1:**
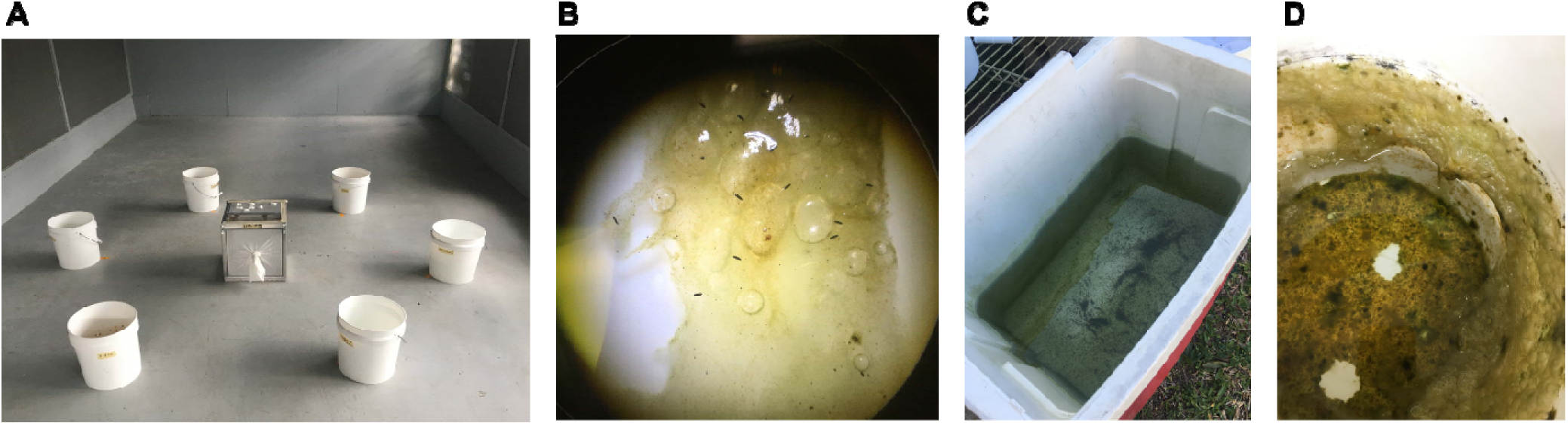
Photographs of periphyton specimens from a field-collected container and experimental buckets taken during an oviposition choice experiment at a semi-field cage in the CDC dengue branch campus, San Juan, Puerto Rico. **(A)** Photograph of semi-field cage set-up during one experimental replicate. Three periphyton-containing buckets and three control buckets were randomly placed at equidistant points around a cage of gravid mosquitoes, which were released into the field cage and allowed one day to lay eggs. **(B)** Photograph of *Ae. aegypti* eggs laid on periphyton during one replicate of the oviposition choice experiment. Taken via stereo microscope. Photo credits: Luisa Otero-Lopez. **(C)** A photograph of the original container with visible periphyton layer and *Aedes aegypti* immatures, which was collected from a residence in Puerto Nuevo, San Juan, Puerto Rico. Extracts of the periphyton from this container were used to seed the plastic buckets used in the oviposition experiments. **(D)** Photograph of an experimental bucket and periphyton layer taken at 15-weeks after the inoculation with periphyton.

### Egg counting

After mosquitoes were removed, buckets were brought to the laboratory where they were examined for the presence of mosquito eggs (Fig. 1B). Eggs were extracted from both the bucket surface and the water within the bucket. The water from each bucket was poured through white filter paper, which was left to dry on a flat surface. Any periphyton floating in the water, or that ended up on filter papers was inspected for the presence of eggs under a stereo microscope. Most eggs were attached to the periphyton in the container’s walls right above the water line. After removal of the water, the bucket was left to dry for approximately two days, depending on the thickness of the periphyton biofilm. Once dry, masking tape was used to detach and collect all the eggs until no further eggs were visible to the naked eye. Pieces of tape were then gently placed on top of filter paper and egg numbers counted under a stereo microscope. All papers were carefully labelled to match their bucket and treatment of origin.

### Fecundity Data Analysis

To model the number of mosquito eggs laid in response to the different treatments, Gaussian linear mixed-effects model with an identity link function were fitted using the glmmTMB package [31] in R version 4.3.1. In experiment 1, the model included ‘Treatment’ (categorical with two levels: Control and Periphyton) as a fixed effect and ‘Experiment’ as a random intercept to account for the different trials in this experiment. To account for strong heteroscedasticity, treatment-specific residual variances were explicitly modeled using the dispformula argument. In experiment 2, the random intercept was excluded as its variance estimate was effectively zero, and ‘Treatment’ was the sole predictor. Model diagnostics using the DHARMa package [32] confirmed no significant overdispersion or residual violations of assumptions in either experiment.

### Mosquito rearing

For all oviposition experiments we used the Patillas, Puerto Rico strain of *Ae. aegypti*. This strain was originally established in the CDC Dengue Branch insectary in 2012 from eggs collected in ovitraps in the Patillas municipality, located in Southeast Puerto Rico. Post-establishment, field collected mosquitoes from Patillas have been introduced to the colony at least once per year [33, 34]. Mosquitoes were reared under standard insectary conditions: 12:12h (light: dark), 25–27°C, and 75% relative humidity. Larvae were reared at a density of approximately 100 immatures per 1L of dechlorinated water per tray and fed rabbit chow daily (71 - 239 mg per tray per day depending on the developmental stage of the larvae). Upon emergence, adult mosquitoes were fed 10% sucrose solution ad libitum.

### Collection of periphyton and establishment of the culture buckets used in experiments

We collected periphyton from a medium-sized cooler located in the Puerto Nuevo neighborhood, which is part of the San Juan Metropolitan area in Puerto Rico during April 2022. This container was selected based on the visual presence of an established periphyton layer and for the presence of mosquito immatures (Fig. 1C). The container was taken to the Centers for Disease Control and Prevention (CDC) - Dengue branch and left outdoors under partial sunlight for five days, covered with mesh cloth so that any emerging mosquitoes would be contained. During this time, larvae and emerged adults that developed in the container were collected and identified as being *Ae. aegypti* by their morphological traits.

Periphyton from this container was visually verified as being free of mosquito eggs and immatures and then used to seed 36 9L plastic buckets (H 24 cm, D 24 cm). We added 4.5 liters of water to these buckets and left them outside for 24 hours so any chlorine in the water would dissipate. Afterwards, periphyton was scratched off the walls of the cooler, and then 0.5L of this mixture was added to each of the 36 buckets, which were already filled with 4.5L of dechlorinated tap water. 45mg of NPK fertilizer (3-4-3 mg/liter, Dr. Earth Item #703P, Winters, CA, USA) was added to each bucket to help with periphyton establishment [35]. These buckets were left outside around the CDC Dengue Branch campus in San Juan, Puerto Rico in areas that were free of plant canopy and covered with mesh cloth to prevent entry of litter and insects. Each bucket had holes drilled at the 5L level to ensure that excess water due to rainfall would drain, and to promote the formation of a periphyton layer at this level. Buckets were visually inspected 2-3 times per week and if the water level had decreased below the 5L line due to evaporation, dechlorinated water was added to that level. Buckets were left outdoors for 15 weeks, and by the end of this period a visible layer of periphyton had grown (Fig. 1D). During this time, additional buckets were left untreated and upside down (to prevent water from entering) for use as control containers in the oviposition experiments.

### Microbial composition of periphyton specimens

We wanted to understand which microorganisms were consistently present in periphyton specimens as these could potentially be the source of compounds that attract mosquitoes to lay eggs. Periphyton samples were collected from the interior surface of buckets directly into sterile 50mL falcon tubes. 26 different buckets were sampled throughout the course of the 15-week culture period (beginning in April 2022) and during the period when oviposition experiments were conducted (July 21 - August 26, 2022) with a total of 26 periphyton samples collected during these times. Each of these samples was visually inspected to ensure that no mosquito eggs were present on the periphyton as it was collected. Periphyton samples were stored at -20°C immediately after collection. These samples were shipped to MR DNA (Shoalwater, TX, USA) who completed DNA extractions using the DNeasy PowerBiofilm Kit (Qiagen, 24000-50), and then performed PCR amplification, quality assurance, library preparation and Illumina MiSeq sequencing of the V4 region of the 16S rRNA gene with 515F/806R primers to generate 2×300bp paired-end reads. Microbial diversity was delineated at the zero-radius operational taxonomic unit (zOTU) level using a custom database and pipeline [36], meaning that each unique sequence variant could be delineated in the dataset. Raw sequence data for this project is available from the Sequence Read Archive (BioProject ID: PRJNA1374143).

Sequencing data were assessed at multiple taxonomic levels. At all levels, taxa with a maximum abundance of less than 0.1% of total reads in any specimen were not considered in the final analysis. Class-level data were used to identify the most abundant and prevalent bacterial classes across all periphyton specimens. Data at the species level were used to produce abundance plots for the 24 most abundant bacterial species to understand the composition of the different periphyton samples. A further plot was prepared from zOTU level data. In this plot, average abundance was considered as the average percentage of total reads across all samples, and prevalence as the proportion of specimens where the read count for a particular zOTU was greater than zero. Abundance was recorded as a dot plot displaying the mean ± s.e.m., while prevalence was represented as a heat map, with these plots prepared in GraphPad Prism (v10.3.1).

Data at the zOTU level were also used to estimate five alpha diversity metrics. A read count matrix for all zOTUs was produced and analyzed using the Vegan package using R version 4.1.1 in R Studio (Ghost Orchid Release 077589bc for macOS). Simpson and Shannon indices and evenness estimates were calculated using the ‘diversity’ function. The Chao-1 index was calculated using the ‘estimateR’ function. All R scripts are provided in Supplementary File 1. An estimate of microbial richness for the periphyton samples was calculated as the number of zOTUs per sample that accounted for at least 0.1% of reads from that sample. Violin dot plots of these five metrics were prepared in GraphPad Prism (v10.3.1). A co-abundance network map was generated for the 20 most abundant zOTUs across all periphyton samples. Read counts for these zOTUs were used to generate a matrix of spearman coefficients in GraphPad Prism (v10.3.1). From this matrix, we selected all moderately strong correlations between zOTUs (positive correlations of ≥ 0.65 and negative correlations of ≤ -0.65) to use in building the network and identify patterns of co-occurrence between key zOTUs in periphyton specimens. This threshold was selected to exclude minor correlations between taxa, which might be of lower biological importance.

## Results

### *Aedes aegypti* mosquitoes consistently lay more eggs in buckets with a periphyton layer

In experiment one (periphyton vs dechlorinated water), a total of 26,724 eggs were collected across five replicates (Fig. 2A). Mean egg counts (± s.e.m.) from periphyton buckets (1524.0 ± 136.5) were higher than those for control buckets (257.6 ± 39.2). The Gaussian linear mixed-effects model revealed that periphyton treatment was a significant predictor of the number of eggs laid in a bucket (χ² = 85.17, *P* < 0.001) while the random effect due to between-experiment variation had a negligible contribution to the overall variance of the model, suggesting that there were no major differences between the five experiments. Across those experiments, between 361.99% and 639.30% more eggs were laid in buckets containing periphyton than in those containing only dechlorinated water.

**Fig. 2:**
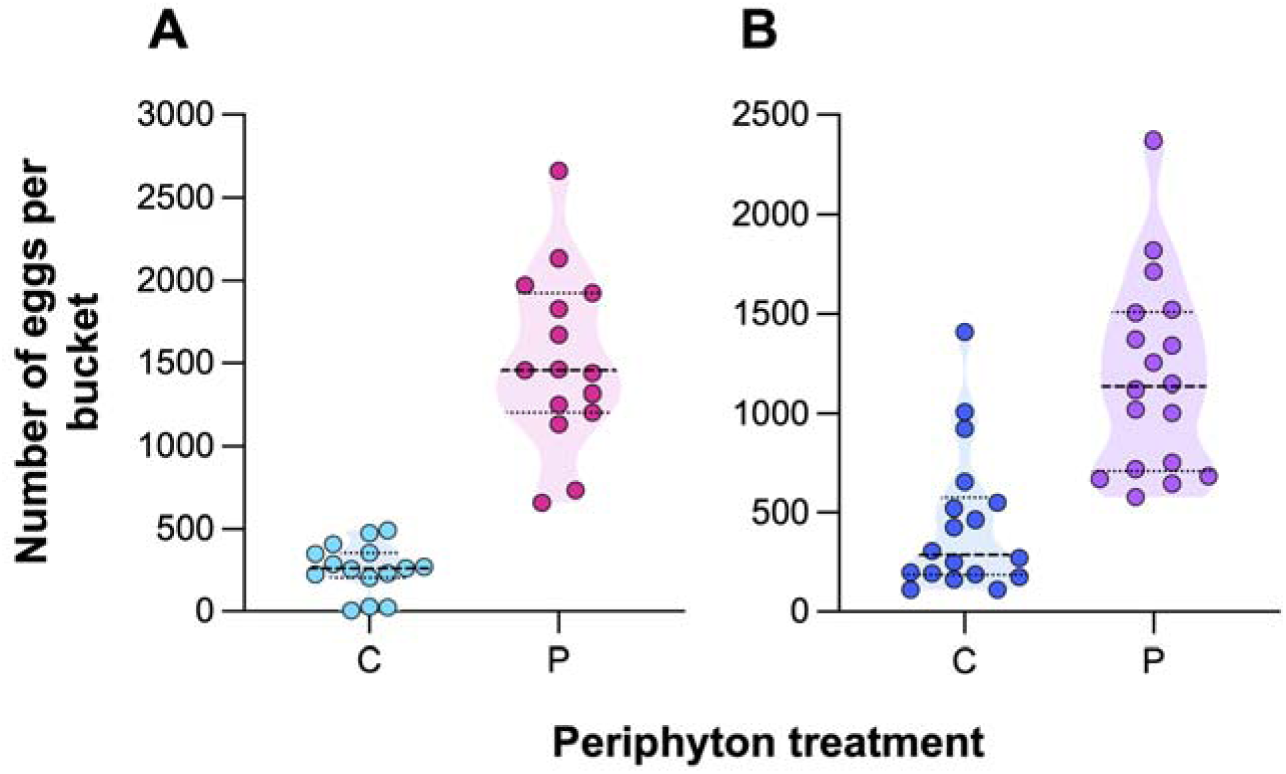
Periphyton containing buckets and water are a highly attractive medium for *Ae. aegypti*. **(A)** Violin dot plots depicting the number of eggs laid by female *Ae. aegypti* in control (C, water only) or periphyton containing buckets (P, periphyton layer on bucket wall) within a semi-field cage. The plot includes all data from five independent replicate oviposition choice assays. **(B)** Violin dot depicting the number of eggs laid by female *Ae. aegypti* in control (C, periphyton in water only) or periphyton containing buckets (P, periphyton layer on bucket wall and in water) within a semi-field cage. The plot includes all data from six independent replicate oviposition choice assays. For both experimental designs, significantly more eggs were laid in the treatment with the greater quantity of periphyton (*P* < 0.0001). Dots represent the total eggs counted from a single bucket. Dashed lines represent treatment medians. Dotted lines represent the 25^th^ and 75^th^ percentiles.

In experiment two (periphyton on bucket walls plus periphyton exposed water vs periphyton exposed water only), a total of 29,172 eggs were collected from buckets across six replicates (Fig. 2B). Mean egg counts (± s.e.m.) from buckets with periphyton on the walls and in the water (1179.9 ± 114.6) were higher than those with periphyton only in the water (440.7 ± 84.5). As with experiment one, the Gaussian linear mixed-effects model revealed that periphyton treatment (χ² = 28.54, *P* < 0.001) was a significant predictor of egg counts in buckets. Across the six experiments, between 28.51% and 487.11% more eggs were laid in buckets with a periphyton layer on the walls than in those without a periphyton layer. Fecundity data are provided in Supplementary File 2, while data for each individual experiment are provided in Supplementary Figure 1.

### Periphyton layers are comprised of diverse microbial communities that include photosynthetic bacteria

A total of 723,145 reads were produced through sequencing of the 16s rRNA gene from periphyton samples. Of these, 718,950 reads were mapped and assigned a matching organism. Periphyton samples produced an average of 27,651.92 reads each, with a minimum of 26,272 and a maximum of 29,022, indicating that all samples had sufficient reads for use in downstream analysis. The average periphyton specimen contained reads associated with Kingdoms Archaea (3.81% of total reads), Bacteria (94.49%), Eukaryota (0.12%), Fungi (0.01%), and Viridiplantae (1.57%). By average percentage of total reads, the five most abundant classes observed were Cyanobacteria (37.89%), Alphaproteobacteria (17.56%), Cytophagia (6.73%), Betaproteobacteria (5.04%), and Planctomycetia (4.08%). Ten of the twenty-four most abundant microbial species in the data set belonged to class Cyanobacteria, with four of those belonging to genus *Leptolyngbya* (Fig. 3A). The top five species by average abundance were *Leptolyngbya frigida* (10.57%), *Oscillatoria amphigranulata* (6.67%), *Chryseolinea* sp. (5.57%), the archaea *Nitrosoarchaeum limnia* (3.87%), and *Phormidium tenue* (3.67%). Analysis of zOTU data revealed several Cyanobacteria were both highly abundant and prevalent (Fig. 3B), including several zOTUs that had the species *Leptolyngbya frigida* as the highest probability match (zOTUs 2, 9, 20, and 23). Similarly, we observed three zOTUs linked to genus *Chryseolinea* (zOTUs 3, 8, and 15) that were also highly abundant and prevalent. Other abundant zOTUs of interest included, *Levilinea saccharolytica* (zOTU4), a non-motile anaerobe, *Calothrix desertica* (zOTU5) a benthic cyanobacterium associated with soils and deserts, and *Rhodopseudomonas pangongensis* (zOTU6), a phototroph. Sequencing data used in this analysis are provided in Supplementary File 3.

**Figure 3:**
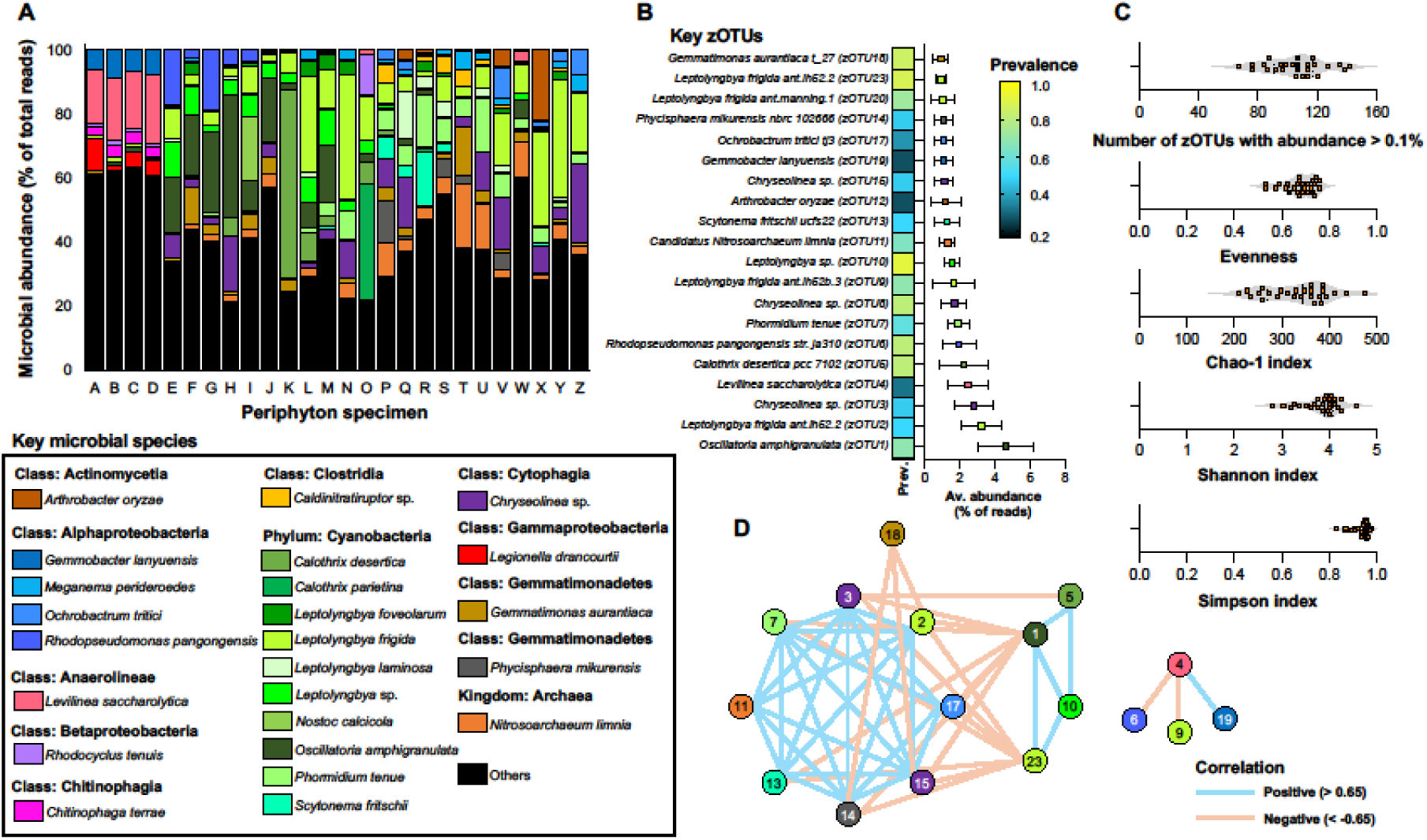
Periphyton samples are comprised of complex microbial communities rich in Cyanobacteria. Microbial profiling of 26 periphyton samples was conducted using 2x 300bp 16s rRNA fingerprinting. **(A)** Stacked bar plot depicting the relative abundance of key microbial taxa found in periphyton samples at the species level, represented as a percentage of total reads per sample. **(B)** Combination heatmap and dot plot depicting the prevalence (proportion of samples where the zOTU was detected) and average abundance (average percentage of total reads across all samples ± s.e.m.) of 20 bacterial zOTUs associated with periphyton samples, indicating that the most common bacteria found in periphyton belonged to classes Alphaproteobacteria and Cytophagia and Phylum Cyanobacteria. Few bacteria had a high average abundance, which suggested that the attractiveness of periphyton to mosquitoes was unlikely caused by a single microorganism. **(C)** Violin dot plots depicting five alpha diversity metrics for periphyton samples: richness (number of zOTUs per sample with abundance > 0.1%), evenness, Chao-1 index, Shannon index, and Simpson index. Each orange dot represents the value for one periphyton sample. These analyses indicate that periphyton specimens have high microbial diversity. **(D)** Network map diagram depicting co-occurrence between key zOTUs in periphyton samples. Only correlations with a strength of > 0.65 (blue) or <-0.65 (orange) are depicted. Positive interactions between taxa might indicate commensal or mutualistic interactions between taxa, while negative interactions might indicate competitive or exclusion interactions. Both of the types of interactions were observed in the periphyton network, suggesting that the community interactions are complex. Numbers in circles correspond to zOTU numbers, as depicted in panel B. Dot color in panels B and D corresponds to the taxonomy/color scheme of microbial species presented in the key at the bottom left of the figure.

Assessment of periphyton sample alpha diversity using five separate metrics, revealed periphyton to be composed of highly diverse microbial communities (Fig. 3C). Mean richness (total number of zOTUs with an abundance at least 0.1%) across all samples was 105.20 (s.e.m. ± 3.78), with a minimum of 67 and a maximum of 142. Mean evenness was 0.67 (s.e.m. ± 0.01), with a minimum of 0.53 and a maximum of 0.76, indicating that zOTUs varied in abundance across specimens. The mean Shannon (3.81 ± 0.08), mean Chao-1 (333.30 ± 13.36) and median Simpson (0.95 ± interquartile range 0.04) values all describe communities with a high level of microbial diversity, meaning that the periphyton community has a high number of species (richness) that are evenly distributed in the community (evenness).

Network analysis of the 20 most abundant zOTUs revealed 52 co-associations with a strength of greater than 0.65 (*N* = 32) or less than -0.65 (*N* = 20) linking 17 different zOTUs (Fig. 3D). This network was split into two elements based on mutual co-associations. The first element was characterized by a suite of positive co-associations between zOTU2 (*Leptolyngbya frigida* ant.lh52.2), zOTU3 (*Chryseolinea* sp.), zOTU7 (*Phormidium tenue*), zOTU11 (Candidatus *Nitrosoarchaeum limnia*), zOTU13 (*Scytonema fritschii* ucfs22), zOTU14 (*Phycisphaera mikurensis* nbrc 102666), zOTU15 (*Chryseolinea* sp.), and zOTU17 (*Ochrobactrum tritici* tj3). That group of zOTUs was then negatively associated with zOTU1 (*Oscillatoria amphigranulata*), zOTU18 (*Gemmatimonas aurantiaca* t_27), and zOTU23 (*Leptolyngbya frigida* ant.lh52.2). Finally, zOTU1, zOTU23, and zOTU5 (*Calothrix desertica* pcc 7102) and zOTU10 (*Leptolyngbya* sp.) all formed a network of positive co-associations. In the second, smaller sub-network, zOTU4 (*Levilinea saccharolytica*) abundance was observed to be negatively correlated to zOTU6 (*Rhodopseudomonas pangongensis* str. ja310) and zOTU9 (*Leptolyngbya frigida* ant.lh52b.3) but positively correlated with zOTU19 (*Gemmobacter lanyuensis*).

## Discussion

Our results highlight that the presence of periphyton in artificial containers has attractive properties for *Ae. aegypti* oviposition, as we observed that significantly higher proportions of the eggs laid in each experiment were associated with periphyton presence. To our knowledge, this is the first time periphyton has been linked to *Ae. aegypti* oviposition preferences. In experiment one, 85.5% of all eggs were laid in periphyton-exposed buckets. In experiment two, 72.8% of all eggs were laid in buckets containing periphyton both on the walls and in the water, highlighting a preference over buckets with periphyton only on the walls. Through 16s rRNA fingerprinting we also demonstrated that periphyton communities used in these experiments are predominantly composed of bacteria, have a high level of complexity and richness, and contain a high abundance of microorganisms belonging to phylum Cyanobacteria, and classes Alphaproteobacteria and Cytophagia, with many of these organisms likely to be photosynthetic or phototrophic. These observations reinforce links between the attractive properties of microorganisms and egg laying behavior in gravid mosquitoes and their potential use for mosquito surveillance and control purposes. Knowing which species of microorganisms attract gravid *Ae. aegypti*, what kairomones these microorganisms release, and what modulates mosquito behavioral responses can be valuable for the future for the development of lures and traps.

Eleven experimental replicates across two different oviposition choice experimental designs revealed that female *Ae. aegypti* consistently preferred to lay their eggs in buckets containing periphyton. These experimental replicates were conducted over a period of 5 weeks, with modest environmental variation taking place across this period. Such environmental variation has been demonstrated to affect flight and oviposition behaviors of mosquitoes during semi-field experiments [11]. We observed minimal replicate-by-replicate variation and that the increased attractiveness of periphyton was consistent across all replicates. In five replicates, the median eggs laid was higher in periphyton buckets than in those containing only water, while in the six replicates of the second experimental design, median eggs laid was higher when buckets had periphyton on the walls and in the water compared to periphyton only in the water.

Oviposition behavior in mosquitoes is a complex trait shaped by visual, olfactory, and physicochemical cues associated with the nature of the container [2] and the water it contains [11], as well as the presence of eukaryotic and prokaryotic organisms [5, 6, 12]. It is not clear what makes periphyton attractive to gravid females, but we consider that its attractive nature is most likely due to the presence of the microorganisms in periphyton or the modulation of physiochemical parameters of the water due to the presence of the periphyton. We had hypothesized that the consistent attraction of mosquitoes to periphyton was due to the presence of one or more microorganisms which were present at high abundance in all periphyton samples that we tested for attractiveness. However, microbial fingerprinting of these samples revealed that the periphyton microbiome could be characterized as being both highly complex and diverse, as revealed by different alpha diversity metrics. Interestingly, few microbial zOTUs were both highly abundant and highly prevalent across all periphyton specimens, which could suggest that multiple zOTUs present in periphyton may produce compounds that attract mosquitoes or prompt oviposition. This is an observation that should be investigated in more depth, as functional redundancies amongst the mosquito microbiota remain poorly understood and could potentially define variable impacts of microbial community composition on mosquito biology [37, 38]. Likewise, functional redundancy in the microbiome could shape the ease with which microbial bio-products or compounds relevant to altering mosquito behavior (repellents and attractants) [39], mosquito mortality (novel insecticidal active ingredients) [40], and vector competence (modulators of mosquito-pathogen interactions) [41] are identified and produced.

Examination of species- and class-level data reveal a pattern of greater consistency across the 26 periphyton specimens, with the most abundant taxa belonging to the Cyanobacteria. Interestingly, we observed that there were both strong positive and strong negative co-associations amongst Cyanobacteria zOTUs, suggesting that some may compete with each other and that they perhaps share biological niches. Microbes from phylum Cyanobacteria have a broad association with mosquitoes, particularly larvae, as they are commonplace across larval mosquito habitats, where they may serve as a food source for larvae [22, 42]. Cyanobacteria have properties that help attract mosquitoes and other organisms and promote egg laying. Cyanobacterial mats that develop in rushes and sawgrass have been positively correlated to the presence of *Anopheles albimanus* larvae [43]. Likewise, biofilms from *Calothrix* sp., which was detected in our periphyton specimens, can act as attractants to freshwater nematodes [44]. Cyanobacteria-mediated increases in temperature and CO_2_ have been speculated to promote oviposition [18]. In addition, the production of the secondary metabolite Geosmin by the Cyanobacteria *Kamptonema* promotes oviposition and attracts the larvae of *Ae. aegypti* [45]. Given that 2-ethyl hexanol and 2,4-di-tert-butylphenol, two volatile organic compounds produced by *Klebsiella* sp., a Gammaproteobacterium, prompt oviposition behavior in *Ae. aegypti* [5] it is likely that mosquitoes are responsive to numerous different cues associated with bacteria and their metabolites. Interestingly, some Cyanobacteria produce endotoxins that can have larvicidal activity [46], while others have failed to demonstrate such impacts [47]. Cyanobacteria have also been modified to express toxins, including those from *Bacillus thuringiensis* [48, 49]. It is unclear whether Cyanobacteria that produce larvicidal endotoxins still promote oviposition in mosquitoes.

Of the other microorganisms that are highly abundant in periphyton, little is known about their interactions with mosquitoes. All three of *Chryseolinea* sp., a clade of bacteria, that are mostly aerobic and soil-associated, *Levilinea saccharolytica,* a filamentous, mesophilic bacterium, and Candidatus *Nitrosoarchaeum limnia*, an Archaeon, have no established links to mosquitoes in the literature. Some members of genus *Leptolyngbya* display minimal larvicidal activity against *Ae. aegypti* [50] while others were part of the Cyanobacteria mats that attract *Anopheles albimanus* [43]. Another key bacterial class in periphyton, Alphaproteobacteria, is commonly found in mosquitoes [51, 52]. It is abundant in several different larval habitats but comparatively less abundant in adult mosquitoes [22]. Interestingly, the microbial composition of periphyton was distinct from that of the water from *Ae. aegypti* tree hole habitats, which had little Cyanobacteria, but did share some Alphaproteobacteria [22]. To better understand the interactions between these abundant microorganisms and *Ae. aegypti* mosquito behavior, future work could examine the attractiveness of these individual bacteria to gravid females seeking an oviposition site.

Given the complexity and diversity of volatile organic compounds produced by microorganisms that stimulate a behavioral response in mosquitoes [53], we were not surprised to observe a preference for oviposition in periphyton-associated buckets in our data. Interestingly, several different plant infusions have been shown to attract gravid *Ae. aegypti* females [33] even though these infusions are prepared with plants materials that are not commonly present in or around *Ae. aegypti* aquatic habitats, like hay infusion made with *Cynodon nlemfuensis* (Vanderyst). Infusions of Bermuda grass, *Cynodon dactylon* (Linnaeus), generate at least 10 compounds due to bacterial catabolism, which can elicit mosquito oviposition activity [11].

Our study possessed several limitations that may have influenced the results. While all of the periphyton specimens we utilized in our study were derived from a single original source, they exhibited variation in microbial composition and physical properties, suggesting that the attractive nature of periphyton to mosquitoes is robust. However, similar investigation of periphyton from other sources would be valuable to assess consistency of attractiveness to mosquitoes as well as microbial composition. It is possible that alternative periphyton communities may have produced a different range of volatile organic compounds and therefore might have differed in their attractiveness to mosquitoes. A yet-to-be investigated parameter in the attraction of periphyton may involve contact cues, which are used by mosquitoes to assess the suitability of a container for egg laying [11]. We observed heterogeneity in the texture of the periphyton layer between buckets, with some having thicker mucilaginous biofilms and others having thinner and coarser biofilms (Fig. 1D). Surfaces with rougher textures [15] and darker colors [54] can help to promote mosquito oviposition behavior. The contribution of such variation to the attractive properties of periphyton is unclear but could have played a role in our experiments. Finally, the physical characteristics of the semi-field cage where our experiments were performed may have influenced mosquito oviposition behavior. It is possible that variation in ventilation, patterns of shade, temperature, humidity, or other physicochemical parameters within the cage may have contributed to the results and to the odor landscape within the cage and/or the periphyton-containing buckets.

## Conclusions

Overall, our data demonstrate that the complex microbial communities that comprise periphyton have attractive properties that should be investigated further and could potentially prove to be valuable in the development of novel lures and traps targeting gravid *Ae. aegypti* females for control purposes. Going forward, it will be interesting to identify specific periphyton-associated microorganisms that mediate attraction in these mosquitoes, and to identify the means through which they do so.

## Supporting information

Supplementary File 1 - R Scripts

Supplementary File 2 - Fecundity experiment data

Supplementary File 3 - Sequencing data

## Figures

**Supplementary Figure 1:**
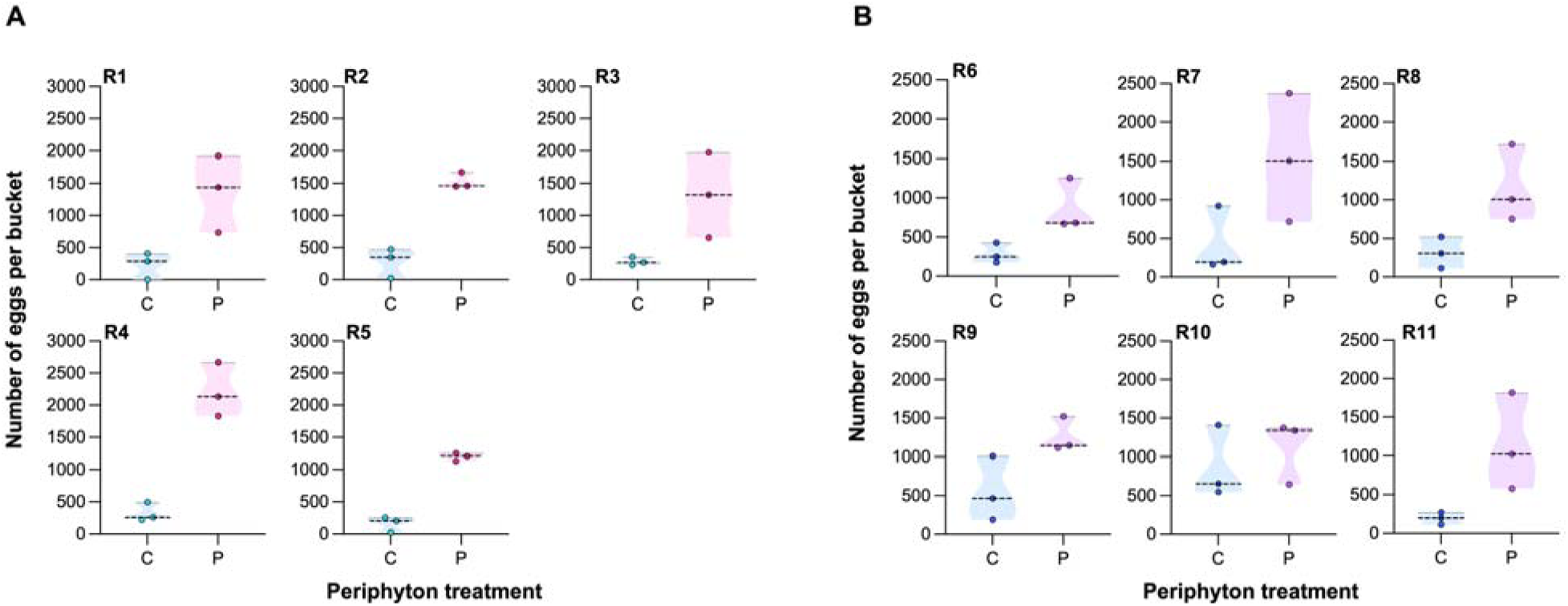
Data from individual fecundity experiments. Violin dot plots for five experimental replicate (R1-R5) oviposition choice assays depicting the number of eggs laid by female *Ae. aegypti* in control (C, water only) or periphyton containing buckets (P, periphyton layer on bucket wall) within a semi-field cage. **(B)** Violin dot plots for six experimental replicate (R6-R11) oviposition choice assays depicting the number of eggs laid by female *Ae. aegypti* in control (C, periphyton in water only) or periphyton containing buckets (P, periphyton layer on bucket wall and in water) within a semi-field cage. Three buckets from each treatment were used in each replicate. Dots represent the total eggs counted from a single bucket. Dashed lines represent treatment medians.

## Abbreviations

CDC: Centers for Disease Control and Prevention
zOTU: zero-radius operational taxonomic unit

## Declarations

### Ethics approval

not applicable

### Consent for publication

not applicable

### Availability of data and materials

All data generated or analysed during this study are included in this published article: Supplementary File 1 contains the script for the Gaussian linear mixed-effects model used for the fecundity data, and all the code for the alpha diversity indices used to analyze the periphyton microbiome data. Oviposition experiments data is available in Supplementary File 2, and the microbial community sequencing data is available in Supplementary File 3. Sequences are also available in the National Institute of Health Sequence Read Archive (SRA) (BioProject ID: PRJNA1374143).

### Authors’ contributions

LMOL conceptualized, visualized, developed the methodology, carried out the field work, the experimental work, wrote the first version of the manuscript and reviewed and edited the final version. AJB performed the fecundity analysis and contributed to the revision of the manuscript’s final version. EPC analyzed the microbial community data, prepared the graphical abstract and drafted the final version of the manuscript. RB conceptualized, visualized, helped develop the methodology and secured funding. All authors have read and agreed to the published version of the manuscript.

## Acknowledgements

the authors thank the field and laboratory technicians that helped during the insectary and experimental work: Jesus Estudillo, Marta Díaz, Keyla Charriez, Orlando Gonzáles, Luis Pérez and Luis Rivera. José Ruiz contributed with useful discussions.

## Disclaimer

The findings and conclusions in this report are those of the authors and do not necessarily represent the views of the US Centers for Disease Control and Prevention.

## References

1. Bentley MD, Day JF. Chemical ecology and behavioral aspects of mosquito oviposition. Annu Rev Entomol. 1989;34:401–21; doi: 10.1146/annurev.en.34.010189.002153. https://www.ncbi.nlm.nih.gov/pubmed/2564759.

2. David MR, Maciel-de-Freitas R, Petersen MT, Bray D, Hawkes FM, Fernandez-Grandon GM, et al. *Aedes aegypti* oviposition-sites choice under semi-field conditions. Med Vet Entomol. 2023;37 4:683–92; doi: 10.1111/mve.12670. https://www.ncbi.nlm.nih.gov/pubmed/37265439.

3. Harrington LC, Ponlawat A, Edman JD, Scott TW, Vermeylen F. Influence of container size, location, and time of day on oviposition patterns of the dengue vector, *Aedes aegypti*, in Thailand. Vector Borne Zoonotic Dis. 2008;8 3:415–23; doi: 10.1089/vbz.2007.0203. https://www.ncbi.nlm.nih.gov/pubmed/18279006.

4. Vezzani D, Albicocco AP. The effect of shade on the container index and pupal productivity of the mosquitoes *Aedes aegypti* and *Culex pipiens* breeding in artificial containers. Med Vet Entomol. 2009;23 1:78–84; doi: 10.1111/j.1365-2915.2008.00783.x. https://www.ncbi.nlm.nih.gov/pubmed/19239617.

5. Mosquera KD, Khan Z, Wondwosen B, Alsanius B, Hill SR, Ignell R, et al. Odor-mediated response of gravid *Aedes aegypti* to mosquito-associated symbiotic bacteria. Acta Trop. 2023;237:106730; doi: 10.1016/j.actatropica.2022.106730. https://www.ncbi.nlm.nih.gov/pubmed/36280207.

6. Ponnusamy L, Wesson DM, Arellano C, Schal C, Apperson CS. Species composition of bacterial communities influences attraction of mosquitoes to experimental plant infusions. Microb Ecol. 2010;59 1:158–73; doi: 10.1007/s00248-009-9565-1. https://www.ncbi.nlm.nih.gov/pubmed/19641948.

7. Ponnusamy L, Xu N, Nojima S, Wesson DM, Schal C, Apperson CS. Identification of bacteria and bacteria-associated chemical cues that mediate oviposition site preferences by *Aedes aegypti*. Proc Natl Acad Sci U S A. 2008;105 27:9262–7; doi: 10.1073/pnas.0802505105. https://www.ncbi.nlm.nih.gov/pubmed/18607006.

8. Allan SA, Kline DL. Larval rearing water and preexisting eggs influence oviposition by *Aedes aegypti* and *Ae. albopictus* (Diptera: Culicidae). J Med Entomol. 1998;35 6:943–7; doi: 10.1093/jmedent/35.6.943. https://www.ncbi.nlm.nih.gov/pubmed/9835684.

9. Ganesan K, Mendki MJ, Suryanarayana MVS, Prakash S, Malhotra RC. Studies of *Aedes aegypti* (Diptera: Culicidae) ovipositional responses to newly identified semiochemicals from conspecific eggs. Aust J Entomol. 2006;45:75–80; doi: 10.1111/j.1440-6055.2006.00513.x. <GO to ISI>://WOS:000234794300011.

10. Allan SA, Kline DL. Evaluation of organic infusions and synthetic compounds mediating oviposition in *Aedes albopictus* and *Aedes aegypti* (Diptera: Culicidae). J Chem Ecol. 1995;21 11:1847–60; doi: 10.1007/BF02033681. https://www.ncbi.nlm.nih.gov/pubmed/24233834.

11. Day JF. Mosquito Oviposition Behavior and Vector Control. Insects. 2016;7 4; doi: 10.3390/insects7040065. https://www.ncbi.nlm.nih.gov/pubmed/27869724.

12. Zahiri N, Rau ME, Lewis DJ. Starved larvae of *Aedes aegypti* (Diptera: Culicidae) render waters unattractive to ovipositing conspecific females. Environ Entomol. 1997;26 5:1087–90; doi: DOI 10.1093/ee/26.5.1087. <GO to ISI>://WOS:A1997YH73300013.

13. Lowenberger CA, Rau ME. Selective Oviposition by *Aedes aegypti* (Diptera, Culicidae) in Response to a Larval Parasite, *Plagiorchis elegans* (Trematoda, Plagiorchiidae). Environ Entomol. 1994;23 5:1269–76; doi: DOI 10.1093/ee/23.5.1269. <GO to ISI>://WOS:A1994PN32200029.

14. Torres-Estrada JL, Rodriguez MH, Cruz-Lopez L, Arredondo-Jimenez JI. Selective oviposition by *Aedes aegypti* (Diptera: culicidae) in response to *Mesocyclops longisetus* (Copepoda: Cyclopoidea) under laboratory and field conditions. J Med Entomol. 2001;38 2:188–92; doi: 10.1603/0022-2585-38.2.188. https://www.ncbi.nlm.nih.gov/pubmed/11296821.

15. O’Gower AK. Environmental stimuli and the oviposition behaviour of *Aedes aegypti* var. *queenslandis* Theobald (Diptera, Culicidae). Animal Behaviour. 1963;11 1:189–97; doi: 10.1016/0003-3472(63)90028-9.

16. Arbaoui AA, Chua TH. Bacteria as a source of oviposition attractant for *Aedes aegypti* mosquitoes. Trop Biomed. 2014;31 1:134–42. <GO to ISI>://WOS:000334283600015.

17. Chadee DD, Lakhan A, Ramdath WR, Persad RC. Oviposition response of *Aedes aegypti* mosquitoes to different concentrations of hay infusion in Trinidad, West Indies. J Am Mosq Control Assoc. 1993;9 3:346–8. https://www.ncbi.nlm.nih.gov/pubmed/8245947.

18. Girard M, Martin E, Vallon L, Raquin V, Bellet C, Rozier Y, et al. Microorganisms Associated with Mosquito Oviposition Sites: Implications for Habitat Selection and Insect Life Histories. Microorganisms. 2021;9 8; doi: 10.3390/microorganisms9081589. https://www.ncbi.nlm.nih.gov/pubmed/34442667.

19. Ponnusamy L, Xu N, Boroczky K, Wesson DM, Abu Ayyash L, Schal C, et al. Oviposition responses of the mosquitoes *Aedes aegypti* and *Aedes albopictus* to experimental plant infusions in laboratory bioassays. J Chem Ecol. 2010;36 7:709–19; doi: 10.1007/s10886-010-9806-2. https://www.ncbi.nlm.nih.gov/pubmed/20521087.

20. Reiter P, Amador MA, Colon N. Enhancement of the CDC ovitrap with hay infusions for daily monitoring of *Aedes aegypti* populations. J Am Mosq Control Assoc. 1991;7 1:52–5. https://www.ncbi.nlm.nih.gov/pubmed/2045808.

21. Muturi EJ, Orindi BO, Kim CH. Effect of leaf type and pesticide exposure on abundance of bacterial taxa in mosquito larval habitats. PLoS One. 2013;8 8:e71812; doi: 10.1371/journal.pone.0071812. https://www.ncbi.nlm.nih.gov/pubmed/23940789.

22. Caragata EP, Otero LM, Tikhe CV, Barrera R, Dimopoulos G. Microbial Diversity of Adult *Aedes aegypti* and Water Collected from Different Mosquito Aquatic Habitats in Puerto Rico. Microb Ecol. 2022;83 1:182–201; doi: 10.1007/s00248-021-01743-6. https://www.ncbi.nlm.nih.gov/pubmed/33860847.

23. Barrera R, Acevedo-Soto V, Ruiz-Valcarcel J, Medina J, Rivera R, Otero L, et al. Defining *Aedes aegypti* density thresholds for preventing human arboviral infections. Acta Trop. 2025;267:107688; doi: 10.1016/j.actatropica.2025.107688. https://www.ncbi.nlm.nih.gov/pubmed/40480602.

24. Barrera R, Amador M, Acevedo V, Hemme RR, Felix G. Sustained, area-wide control of *Aedes aegypti* using CDC autocidal gravid ovitraps. Am J Trop Med Hyg. 2014;91 6:1269–76; doi: 10.4269/ajtmh.14-0426. https://www.ncbi.nlm.nih.gov/pubmed/25223937.

25. Barrera R, Harris A, Hemme RR, Felix G, Nazario N, Munoz-Jordan JL, et al. Citywide Control of *Aedes aegypti* (Diptera: Culicidae) during the 2016 Zika Epidemic by Integrating Community Awareness, Education, Source Reduction, Larvicides, and Mass Mosquito Trapping. J Med Entomol. 2019;56 4:1033–46; doi: 10.1093/jme/tjz009. https://www.ncbi.nlm.nih.gov/pubmed/30753539.

26. Larned ST. A prospectus for periphyton: recent and future ecological research. J N Am Benthol Soc. 2010;29 1:182–206; doi: 10.1899/08-063.1. <GO to ISI>://WOS:000275023500011.

27. Herren CM, Webert KC, McMahon KD. Environmental Disturbances Decrease the Variability of Microbial Populations within Periphyton. mSystems. 2016;1 3; doi: 10.1128/mSystems.00013-16. https://www.ncbi.nlm.nih.gov/pubmed/27822539.

28. Moschini-Carlos V, Henry R, Pompêo MLM. Seasonal variation of biomass and productivity of the periphytic community on artificial substrata in the Jurumirim Reservoir (Sao Paulo, Brazil). Hydrobiologia. 2000;434 1–3:35-40; doi: DOI 10.1023/A:1004086623922. <GO to ISI>://WOS:000089982800004.

29. Roos PJ. Dynamics of periphytic communities. In: Wetzel RG, editor. Periphyton of Freshwater Ecosystems: Proceedings of the First International Workshop on Periphyton of Freshwater Ecosystems held in Växjö, Sweden 14-17 September 1982, 1 edn. The Netherlands: Springer Dordrecht; 1983. p. 356.

30. van Dijk GM. Dynamics and Attenuation Characteristics of Periphyton Upon Artificial Substratum under Various Light Conditions and Some Additional Observations on Periphyton Upon *Potamogeton pectinatus* L. Hydrobiologia. 1993;252 2:143–61; doi: Doi 10.1007/Bf00008152. <GO to ISI>://WOS:A1993KY41900003.

31. Brooks ME, Kristensen K, van Benthem KJ, Magnusson A, Berg CW, Nielsen A, et al. glmmTMB Balances Speed and Flexibility Among Packages for Zero-inflated Generalized Linear Mixed Modeling. R J. 2017;9 2:378–400; doi: Doi 10.32614/Rj-2017-066. <GO to ISI>://WOS:000423751200026.

32. Hartig F: DHARMa: Residual Diagnostics for Hierarchical (Multi-Level / Mixed) Regression Models. 2024.

33. Mackay AJ, Amador M, Barrera R. An improved autocidal gravid ovitrap for the control and surveillance of *Aedes aegypti*. Parasit Vectors. 2013;6 1:225; doi: 10.1186/1756-3305-6-225. https://www.ncbi.nlm.nih.gov/pubmed/23919568.

34. Poole-Smith BK, Hemme RR, Delorey M, Felix G, Gonzalez AL, Amador M, et al. Comparison of vector competence of *Aedes mediovittatus* and *Aedes aegypti* for dengue virus: implications for dengue control in the Caribbean. PLoS Negl Trop Dis. 2015;9 2:e0003462; doi: 10.1371/journal.pntd.0003462. https://www.ncbi.nlm.nih.gov/pubmed/25658951.

35. Darriet F. Synergistic Effect of Fertilizer and Plant Material Combinations on the Development of *Aedes aegypti* (Diptera: Culicidae) and *Anopheles gambiae* (Diptera: Culicidae) Mosquitoes. J Med Entomol. 2018;55 2:496–500; doi: 10.1093/jme/tjx231. https://www.ncbi.nlm.nih.gov/pubmed/29309617.

36. Glassing A, Dowd SE, Galandiuk S, Davis B, Jorden JR, Chiodini RJ. Changes in 16s RNA Gene Microbial Community Profiling by Concentration of Prokaryotic DNA. J Microbiol Methods. 2015;119:239–42; doi: 10.1016/j.mimet.2015.11.001. https://www.ncbi.nlm.nih.gov/pubmed/26569458.

37. Li L, Wang T, Ning Z, Zhang X, Butcher J, Serrana JM, et al. Revealing proteome-level functional redundancy in the human gut microbiome using ultra-deep metaproteomics. Nat Commun. 2023;14 1:3428; doi: 10.1038/s41467-023-39149-2. https://www.ncbi.nlm.nih.gov/pubmed/37301875.

38. Tian L, Wang XW, Wu AK, Fan Y, Friedman J, Dahlin A, et al. Deciphering functional redundancy in the human microbiome. Nat Commun. 2020;11 1:6217; doi: 10.1038/s41467-020-19940-1. https://www.ncbi.nlm.nih.gov/pubmed/33277504.

39. Kajla MK, Barrett-Wilt GA, Paskewitz SM. Bacteria: A novel source for potent mosquito feeding-deterrents. Sci Adv. 2019;5 1:eaau6141; doi: 10.1126/sciadv.aau6141. https://www.ncbi.nlm.nih.gov/pubmed/30746455.

40. Caragata EP, Otero LM, Carlson JS, Borhani Dizaji N, Dimopoulos G. A Nonlive Preparation of *Chromobacterium* sp. Panama (Csp_P) Is a Highly Effective Larval Mosquito Biopesticide. Appl Environ Microbiol. 2020;86 11; doi: 10.1128/AEM.00240-20. https://www.ncbi.nlm.nih.gov/pubmed/32220845.

41. Zhang L, Wang D, Shi P, Li J, Niu J, Chen J, et al. A naturally isolated symbiotic bacterium suppresses flavivirus transmission by *Aedes* mosquitoes. Science. 2024;384 6693:eadn9524; doi: 10.1126/science.adn9524. https://www.ncbi.nlm.nih.gov/pubmed/38669573.

42. Thiery I, Nicolas L, Rippka R, Tandeau de Marsac N. Selection of cyanobacteria isolated from mosquito breeding sites as a potential food source for mosquito larvae. Appl Environ Microbiol. 1991;57 5:1354–9; doi: 10.1128/aem.57.5.1354-1359.1991. https://www.ncbi.nlm.nih.gov/pubmed/1677241.

43. Rejmankova E, Roberts DR, Manguin S, Pope KO, Komarek J, Post RA. *Anopheles albimanus* (Diptera: Culicidae) and cyanobacteria: An example of larval habitat selection. Environ Entomol. 1996;25 5:1058–67; doi: DOI 10.1093/ee/25.5.1058. <GO to ISI>://WOS:A1996VR84800024.

44. Höckelmann C, Moens T, Jüttner F. Odor compounds from cyanobacterial biofilms acting as attractants and repellents for free-living nematodes. Limnol Oceanogr. 2004;49 5:1809–19; doi: DOI 10.4319/lo.2004.49.5.1809. <GO to ISI>://WOS:000224979900032.

45. Melo N, Wolff GH, Costa-da-Silva AL, Arribas R, Triana MF, Gugger M, et al. Geosmin Attracts *Aedes aegypti* Mosquitoes to Oviposition Sites. Curr Biol. 2020;30 1:127–34 e5; doi: 10.1016/j.cub.2019.11.002. https://www.ncbi.nlm.nih.gov/pubmed/31839454.

46. Kiviranta J, Abdelhameed A, Sivonen K, Niemela SI, Carlberg G. Toxicity of Cyanobacteria to Mosquito Larvae Screening of Active Compounds. Environ Toxic Water. 1993;8 1:63–71; doi: DOI 10.1002/tox.2530080107. <GO to ISI>://WOS:A1993KH61100006.

47. Vazquez-Martinez MG, Rodriguez MH, Arredondo-Jimenez JI, Mendez-Sanchez JD, Bond-Compean JG, Cold-Morgan M. Cyanobacteria associated with *Anopheles albimanus* (Diptera: Culicidae) larval habitats in southern Mexico. J Med Entomol. 2002;39 6:825–32; doi: 10.1603/0022-2585-39.6.825. https://www.ncbi.nlm.nih.gov/pubmed/12495179.

48. Boussiba S, Wu XQ, Ben-Dov E, Zarka A, Zaritsky A. Nitrogen-fixing cyanobacteria as gene delivery system for expressing mosquitocidal toxins of *Bacillus thuringiensis* ssp. *israelensis*. J Appl Phycol. 2000;12 3–5:461-7; doi: Doi 10.1023/A:1008114929490. <GO to ISI>://WOS:000089987100034.

49. Ketseoglou I, Bouwer G. The persistence and ecological impacts of a cyanobacterium genetically engineered to express mosquitocidal *Bacillus thuringiensis* toxins. Parasit Vectors. 2016;9 1:273; doi: 10.1186/s13071-016-1544-z. https://www.ncbi.nlm.nih.gov/pubmed/27165108.

50. Ayala FI, Becerra LM, Quintana J, Bayona LM, Ramos FA, Puyana M, et al. Environmental and cultured cyanobacteria as sources of *Aedes aegypti* larvicides. Universitas Scientiarum. 2019;24 3:465–96; doi: 10.11144/Javeriana.SC24-3.eacc.

51. Buck M, Nilsson LK, Brunius C, Dabire RK, Hopkins R, Terenius O. Bacterial associations reveal spatial population dynamics in *Anopheles gambiae* mosquitoes. Sci Rep. 2016;6:22806; doi: 10.1038/srep22806. https://www.ncbi.nlm.nih.gov/pubmed/26960555.

52. Coon KL, Brown MR, Strand MR. Mosquitoes host communities of bacteria that are essential for development but vary greatly between local habitats. Mol Ecol. 2016;25 22:5806–26; doi: 10.1111/mec.13877. https://www.ncbi.nlm.nih.gov/pubmed/27718295.

53. Coutinho-Abreu IV, Jamshidi O, Raban R, Atabakhsh K, Merriman JA, Akbari OS. Identification of human skin microbiome odorants that manipulate mosquito landing behavior. Sci Rep. 2024;14 1:1631; doi: 10.1038/s41598-023-50182-5. https://www.ncbi.nlm.nih.gov/pubmed/38238397.

54. Muir LE, Kay BH, Thorne MJ. *Aedes aegypti* (Diptera: Culicidae) vision: response to stimuli from the optical environment. J Med Entomol. 1992;29 3:445–50; doi: 10.1093/jmedent/29.3.445. https://www.ncbi.nlm.nih.gov/pubmed/1625292.

