## Supplementary File 1 - R Scripts for "The role of container periphyton as an oviposition attractant for female *Aedes aegypti*"

**Part 1: The code used to analyze the fecundity data is listed below.**

\# Manage file paths for project directory

library(here)

\# Data import

library(readxl)

\# Data wrangling & plotting

library(tidyverse)

library(ggsci)	# Optional: Color palettes

library(patchwork)	# Optional: Plot combination

\# Modeling & diagnostics

library(glmmTMB)

library(DHARMa)

library(car)	# For ANOVA tests

library(emmeans)	# Estimated marginal means

library(multcomp)	# Post hoc comparisons

library(multcompView) # Letter display for group differences

\# Tidying & reporting

library(broom.mixed)

library(easystats)	# Model checks and stats summaries

\# Define folder structure

data_folder <- "Data"

plot_folder <- "Plots"

here <- here::here

\# Experiment 1 ----

\# Load data

exp_1 <- read_excel(here(data_folder, "Exp 1.xlsx"))

\## Final model fit and analysis ----

\# Final model: Linear mixed model with heterogeneous variance across treatments

mod_var <- glmmTMB(

Eggs ~ Treatment + (1 | Experiment),

dispformula = ~ Treatment, # Model treatment-specific variance

family = gaussian,

data = exp_1

)

\# Model diagnostics: Simulated residuals

simulateResiduals(mod_var, plot = TRUE)

\# Significance of fixed effects

Anova(mod_var) # Type II Wald Chi-square test

\# Estimated marginal means and group comparisons

emm <- emmeans(mod_var, pairwise ~ Treatment)

\# Compact letter display for grouping significance

mod_emm <- emmeans(mod_var, ~ Treatment) |>

multcomp::cld(Letters = "ABCD", alpha = 0.05)

\## Visualizations ----

\# (a) Boxplot of Raw Data

BoxP <- ggplot(exp_1, aes(x = Treatment, y = Eggs, fill = Treatment)) +

geom_boxplot(show.legend = FALSE) +

geom_point(position = position_jitterdodge(jitter.width = 0.5, jitter.height = 0.1), alpha = 0.25, size = 2, show.legend = FALSE) +

ylim(0, 2700) +

scale_fill_nejm() +

labs(x = "Treatment", y = "Egg Count") +

theme_modern()

\# (b) Bar Plot: Estimated Means with Error Bars and Letters

BarP <- ggplot(mod_emm, aes(x = Treatment, y = emmean, fill = Treatment)) +

geom_col(color = "black", width = 0.7, show.legend = FALSE) +

geom_errorbar(aes(ymin = lower.CL, ymax = upper.CL), width = 0.1) +

ylim(0, 2700) +

geom_text(aes(label = .group, y = upper.CL + 60), size = 5, vjust = 0.1) +

scale_fill_nejm() +

labs(x = "Treatment", y = "Egg Count ± 95% CI") +

theme_modern()

\# (c) Combined Plot

plots_exp1 <- (BoxP + BarP) + plot_annotation(

title = "Experiment 1 Results",

tag_levels = 'A'

)

plots_exp1

\# Experiment 2 ----

\# Load data

exp_2 <- read_excel(here(data_folder, "Exp 2.xlsx"))

\## Final model fit and analysis ----

\# Final model: Simpler fixed-effect Gaussian model (no random effect needed)

mod_basic <- glmmTMB(

Eggs ~ Treatment,

family = gaussian,

data = exp_2

)

\# Model diagnostics: Simulated residuals

simulateResiduals(mod_basic, plot = TRUE)

\# Significance of fixed effects

Anova(mod_basic)

\# Estimated marginal means and group comparisons

emm <- emmeans(mod_basic, pairwise ~ Treatment)

\# Compact letter display for grouping significance

mod_emm <- emmeans(mod_basic, ~ Treatment) %>%

multcomp::cld(Letters = "ABCD", alpha = 0.05)

\## Visualizations ----

\# (a) Boxplot of Raw Data

BoxP <- ggplot(exp_2, aes(x = Treatment, y = Eggs, fill = Treatment)) +

geom_boxplot(show.legend = FALSE) +

geom_point(position = position_jitterdodge(jitter.width = 0.5, jitter.height = 0.1), alpha = 0.25, size = 2, show.legend = FALSE) +

ylim(0, 2400) +

scale_fill_nejm() +

labs(x = "Treatment", y = "Egg Count") +

theme_modern()

\# (b) Bar Plot: Estimated Means with Error Bars and Letters

BarP <- ggplot(mod_emm, aes(x = Treatment, y = emmean, fill = Treatment)) +

geom_col(color = "black", width = 0.7, show.legend = FALSE) +

geom_errorbar(aes(ymin = lower.CL, ymax = upper.CL), width = 0.1) +

ylim(0, 2400) +

geom_text(aes(label = .group, y = upper.CL + 50), size = 5, vjust = 0.1) +

scale_fill_nejm() +

labs(x = "Treatment", y = "Egg Count ± 95% CI") +

theme_modern()

#Combined Plot

plots_exp2 <- (BoxP + BarP) + plot_annotation(

title = "Experiment 2 Results",

tag_levels = 'A'

)

plots_exp2

**Part 2: The code for the alpha diversity indices used to analyze the periphyton microbiome data is listed below.**

library(vegan)

##PCM – **P**eriphyton **C**ount **M**atrix – a count matrix of zOTU-level data##

#Shannon Index

shannon<-diversity(PCM,index = "shannon")

shannon

plot(shannon)

#Simpson Index

simpson<-diversity(PCM,index = "simpson")

simpson

plot(simpson)

#Chao's Species Estimator

data_richness <- estimateR(PCM)

data_richness

plot(data_richness)

#Eveness Index

data_eveness <- diversity(PCM) / log(specnumber(PCM))

data_eveness

plot(data_eveness)
